# Variation in genetic relatedness is determined by the aggregate recombination process

**DOI:** 10.1101/2020.05.25.115048

**Authors:** Carl Veller, Nathaniel B. Edelman, Pavitra Muralidhar, Martin A. Nowak

**Affiliations:** Department of Organismic and Evolutionary Biology, Harvard University, Massachusetts, USA; Department of Mathematics, Harvard University, Massachusetts, USA

## Abstract

The genomic proportion that two relatives share identically by descent—their genetic relatedness— can vary depending on the history of recombination and segregation in their pedigree. This variation is important in many applications of genetics, including pedigree-based estimation of the genetic variance and heritability of traits, and estimation of pedigree relationships from sequence data. Here, we calculate the variance of genetic relatedness for general pedigree relationships, making no assumptions about the recombination process. For the specific relationships of grandparent-grandoffspring and siblings, the variance of genetic relatedness is a simple decreasing function of 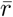, the average proportion of locus pairs that recombine in meiosis. For general pedigree relationships, the variance of genetic relatedness is likewise the average of some function of pairwise recombination rates. Therefore, features of the aggregate recombination process that affect 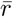 and analogs also affect variance in genetic relatedness. Such features include the number of chromosomes and heterogeneity in their size, and the number of crossovers and their location along chromosomes. Our calculations help to explain several recent observations about variance in genetic relatedness, including that it is reduced by crossover interference (which is known to increase 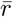). Our methods further allow us to calculate the neutral variance of ancestry among F2s in a hybrid cross, enabling precise statistical inference in F2-based tests for various kinds of selection.

## 1 Introduction

Variance in the amount of DNA shared by relatives identically by descent (IBD)—variance in genetic relatedness—is an important quantity in genetics (Thompson 2013). It translates to variance in the phenotypic similarity of relatives, and is a vital component of pedigree-based estimates of heritability and the genetic variance of traits (Visscher et al. 2006, 2007; Young et al. 2018). It is also an important consideration when estimating pedigree relationships and the degree of inbreeding from genotype data (Kardos et al. 2015; Wang 2016). Variance in genetic relatedness has also been hypothesized to have important evolutionary consequences, including driving the evolution of karyotypes and recombination rates in some clades (Sherman 1979; Wilfert et al. 2007). Moreover, as we show elsewhere, variance in genetic relatedness plays a key role in selection against deleterious introgressed DNA following hybridization (Veller et al. 2019a).

For most pedigree relationships, genetic relatedness can vary because of variable patterns of recombination and segregation within the pedigree. For example, it is possible that a mother segregates only crossoverless paternal chromatids to an egg, in which case the resulting offspring inherits one half of its genome from its maternal grandfather and none from its maternal grandmother. On the other hand, if the mother shuffles her maternal and paternal DNA thoroughly into the egg, the offspring will be approximately equally related (genetically) to its maternal grandparents. Thus, intuitively, a higher degree of genetic shuffling within a pedigree leads to lower variance in genetic relatedness between relatives.

In theoretical calculations of the variance of genetic relatedness, it has typically been assumed that recombination is uniform along chromosomes and that crossover interference is absent [e.g., Franklin (1977); Hill (1993b); Guo (1996); Visscher et al. (2006); a general treatment under this assumption is given by Hill and Weir (2011)]. White and Hill (2020) have recently developed a procedure to estimate the variance of genetic relatedness from linkage maps, without the assumption of uniform recombination rates. However, their method assumes uniform recombination rates in regions between markers (restricting the method to high-density linkage maps) and ignores crossover interference.

In this paper, we derive a general, assumption-free formulation for the variance of genetic relatedness in terms of aggregate genetic shuffling. We demonstrate that the variance of genetic relatedness is a simple, decreasing function of certain newly-developed metrics of genome-wide genetic shuffling: 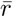 and analogs (Veller et al. 2019b). These metrics, in a natural and intuitive way, take into account features of the aggregate recombination process such as the number of chromosomes and differences in their size, the number of crossovers and their location along the chromosomes, and the spatial relations of crossovers with respect to each other (crossover interference). Many of these features have been overlooked in previous calculations of the variance of genetic relatedness.

Our formulation of the variance of genetic relatedness in terms of 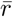 and analogs allows the effect of these meiotic features on the variance of genetic relatedness to be reinterpreted—often with greater intuition—in terms of their effect on aggregate genetic shuffling. For example, it has recently been shown that crossover interference decreases the variance of genetic relatedness (Caballero et al. 2019). This can be explained by the intuitive fact that crossover interference, by spreading crossovers out evenly along chromosomes, increases the amount of genetic shuffling that they cause, thus increasing 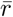 and analogs (Gorlov and Gorlova 2001; Veller et al. 2019b).

In the calculations below, we assume that there is no inbreeding. The number of loci in the genome, *L*, is assumed to be very large, and loci *i* and *j* are recombinant in a random gamete with probability *r*_*ij*_ (e.g., *r*_*ij*_ = 1/2 if *i* and *j* are on different chromosomes). Sex specific recombination rates, 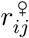 and 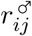, are distinguished where necessary.

## 2 Relationships of direct descent

Pedigree relationships of direct descent (or ‘lineal’ relationships) involve a single lineage, from an ancestor to one of its descendants. We will focus here on the specific example of grandparent-grandoffspring—calculations of the variance of genetic relatedness for general relationships of direct descent are given in SI Section S1.

### Grandparent-grandoffpsring

Let the random variable *IBD*_grand_ be the proportion of a grandoffspring’s genome inherited from a specified grandparent. Consider the gamete produced the grand-offspring’s parent, and let 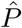 be the fraction of this gamete’s genome that derives from the focal grandparent (so that 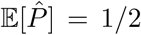). We first wish to calculate 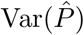. Using an approach similar to that of Hill (1993a) and Visscher et al. (2006), but without making any assumptions about the recombination process, we calculate (details in SI Section S1) that

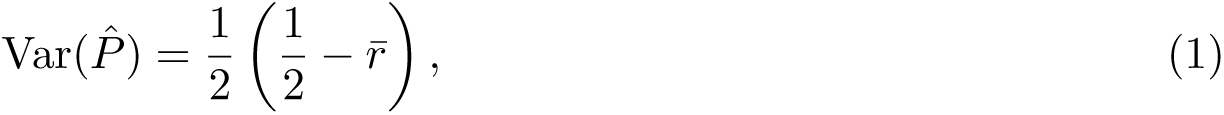

where 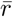 is the probability that a randomly chosen locus pair recombines in meiosis (Veller et al. 2019b). Because half of the grandoffspring’s genome comes from this gamete, 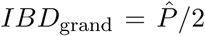, so that 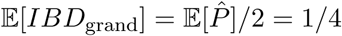 is the coefficient of relationship, and

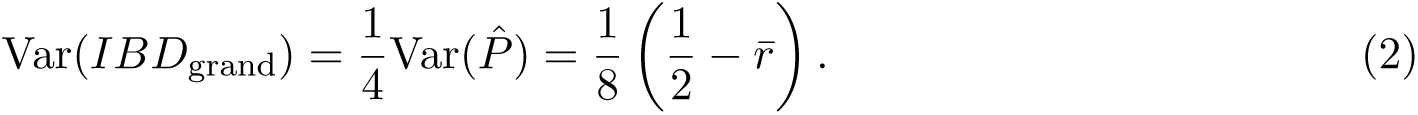

A graphical demonstration of Eq. (2), based on the possible segregation patterns of a given parental meiosis, is shown in Fig. 1. The population variance can be seen as an average across all such meioses. Note that the formulation in Eq. (2) and other such formulations in this paper apply to the whole genome, or a single chromosome, or any specific genomic region. In the latter cases, 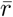 is the probability that a random pair of loci within the region of interest recombine in meiosis. In addition, because the recombination process often differs between the sexes, the value of 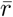 can differ between spermatogenesis and oogenesis. In calculating the variance of genetic relatedness between a grandoffspring and one of its maternal grandparents, the value for oogenesis, 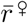, would be used; the value for spermatogenesis, 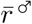, would be used for paternal grandparents.

**Figure 1:**
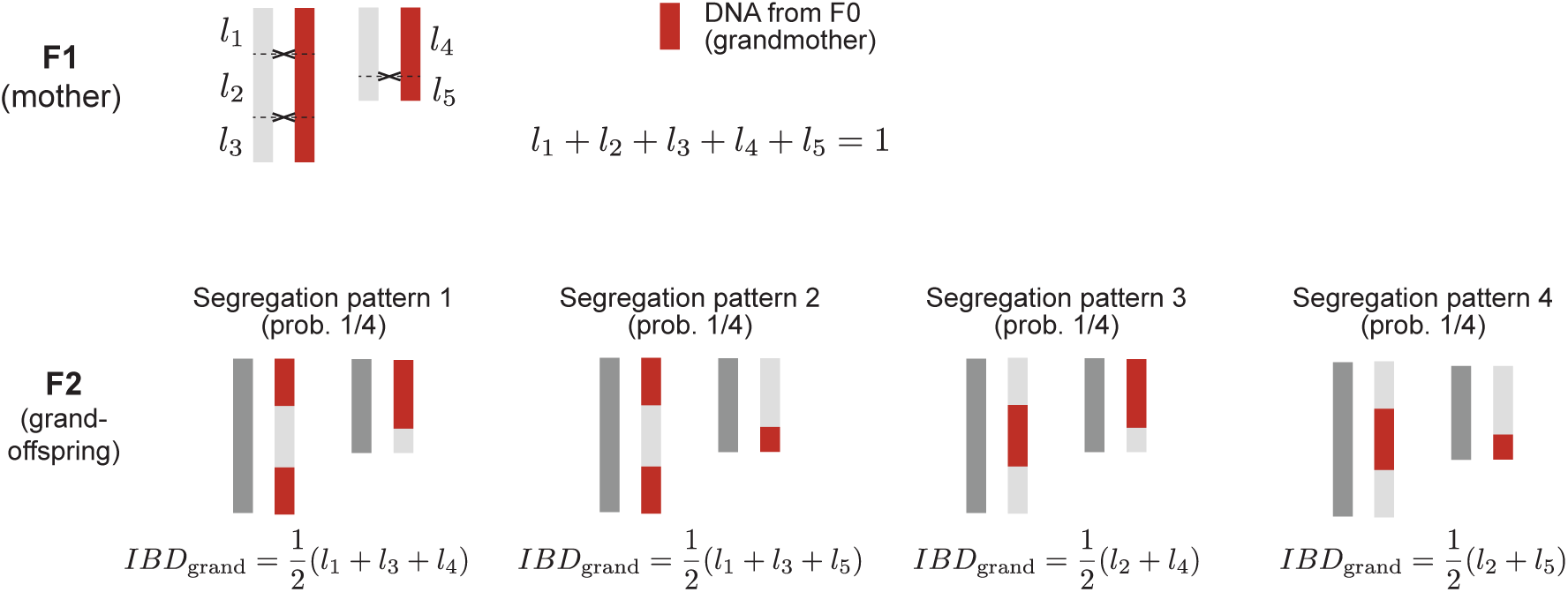
The variance of genetic relatedness between grandoffspring and grandparent, calculated from the possible segregation patterns of a single parental meiosis. In the figure, the positions of crossovers in a maternal meiosis (and the chromatids involved) are specified, but the segregation pattern in the resulting egg (and therefore offspring) is not. Averaging across the four segregation patterns, we find 𝔼[*IBD*_grand_] = (*l*_1_ + *l*_2_ + *l*_3_ + *l*_4_ + *l*_5_)/4 = 1/4, and, from Eq. [1] in Veller et al. (2019b), 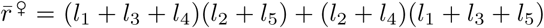. Across the four possible segregation patterns, 𝔼[*IBD*_grand_] = 1/4
and

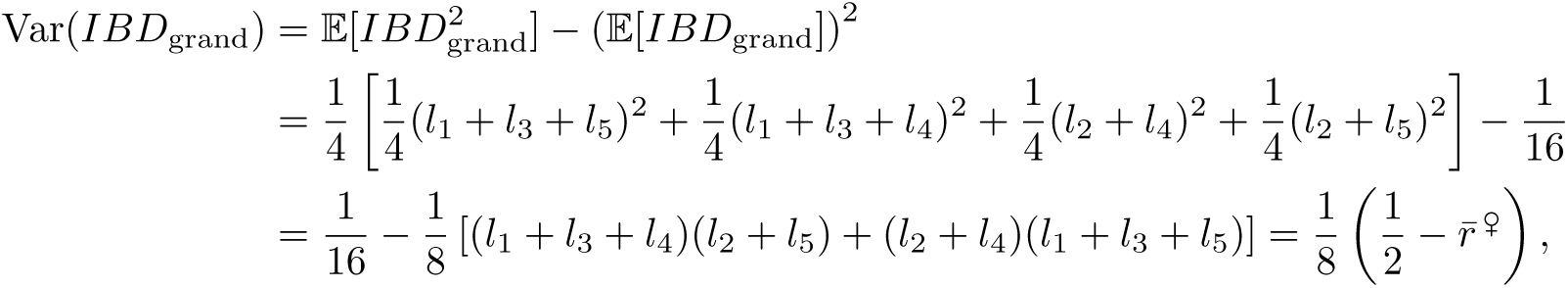

which is Eq. (2).

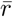 can be estimated from various kinds of data, including cytological data of crossover positions at meiosis I, sequence data from gametes, and linkage maps (Veller et al. 2019b). For example, Veller et al. (2019b) used cytological data from Lian et al. (2008) to estimate an autosomal value for human males of 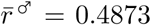. Substituting this value into Eq. (2) reveals that the variance of (autosomal) genetic relatedness of a grandoffspring to its paternal grandparent is Var(*IBD*_grand_) = 1.6 × 10−^3^ (standard deviation *σ* = 0.04, coefficient of variation CV = 16%).

## 3 Indirect relationships

Indirect relationships involve two descendants of at least one individual in the pedigree. In the case of multi-ancestor pedigrees, we will restrict our attention to two-ancestor pedigrees involving a mating between the two ancestors (e.g., full-sibs, aunt-nephew). We focus here on half-siblings and siblings— the calculations for general indirect relationships of this kind are given in SI Section S2.

### Half-siblings

Let the random variable *IBD*_h-sib_ be the proportion of two half-siblings’ genomes that they share IBD, if they have the same father but unrelated mothers. Then 𝔼[*IBD*_h-sib_] = 1/4 is the coefficient of relationship, and

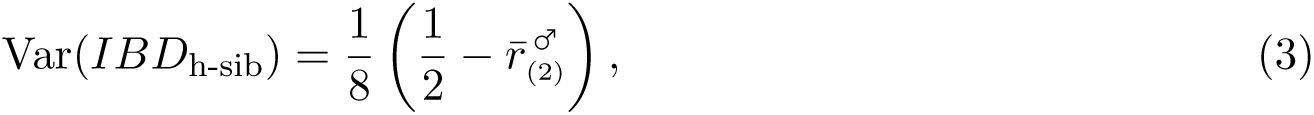

where 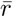 is the probability that a randomly chosen locus pair recombines when the crossovers of two of the father’s meioses are pooled into one meiosis (see Fig. 2 for an example of a pooled meiosis). If the common parent were instead the mother, 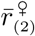 would replace 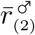. A graphical demonstration of Eq. (3), based on the possible segregation patterns of two meioses in the parent, is given in Fig. 2. Again, the population variance can be seen as an average across all such pairs of meioses.

### Siblings

Let the random variable *IBD*_sib_ be the proportion of two full-siblings’ genomes that they share IBD, assuming their mother and father to be unrelated. Then 𝔼[*IBD*_sib_] = 1/2 is the coefficient of relationship, and

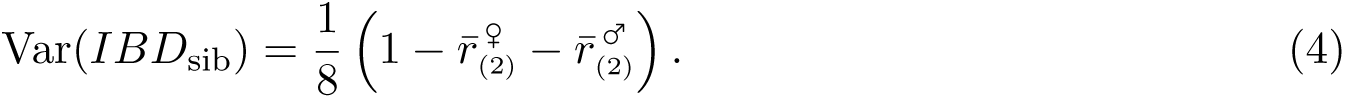

**Figure 2:**
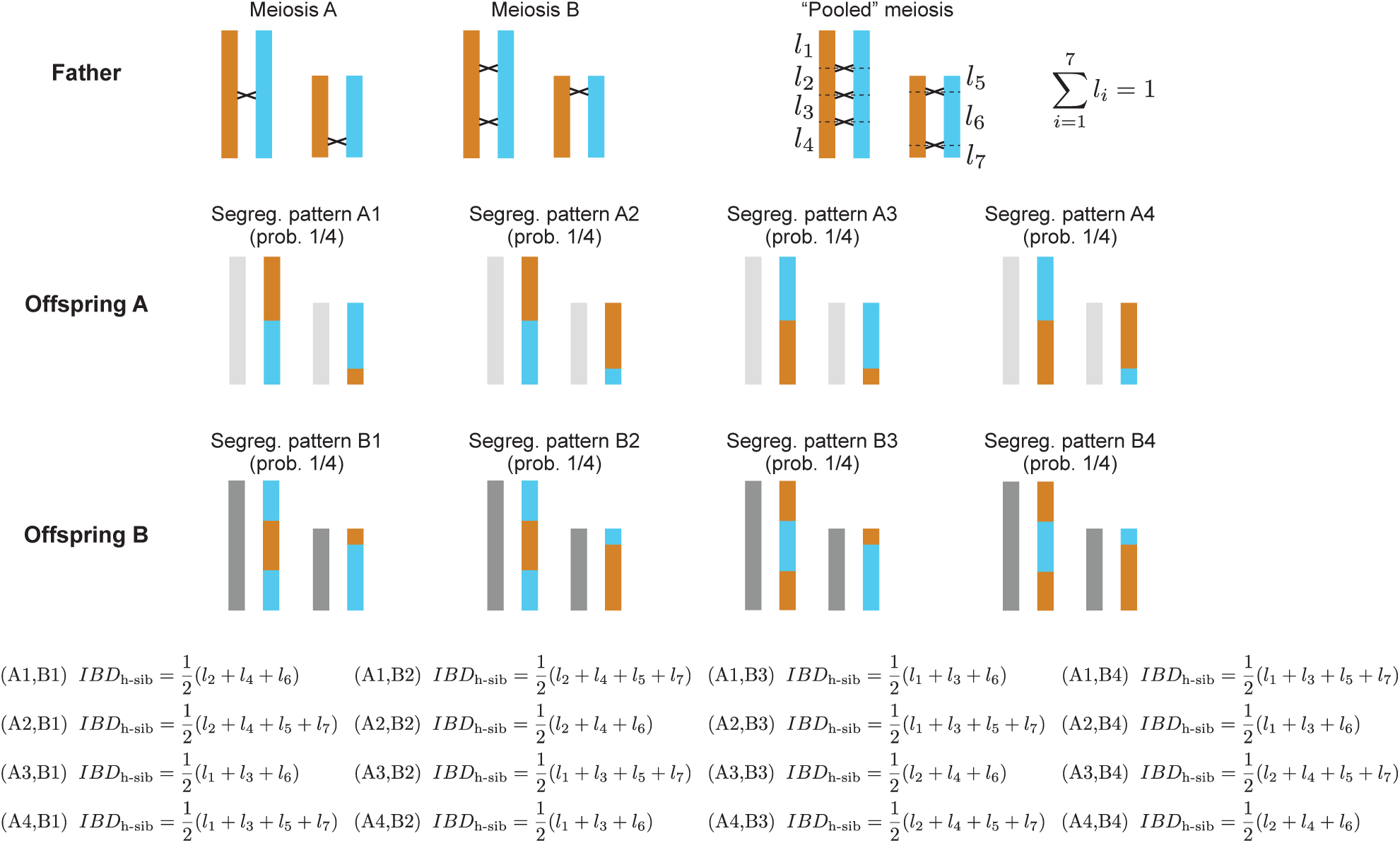
The variance of genetic relatedness between half-siblings, calculated from the possible segregation patterns of two meioses of their common parent. The positions of crossovers in two paternal meioses (and the chromatids involved) are specified, but the segregation patterns in the resulting sperm cells (and therefore the two offspring) are not. Applying Eq. [1] in Veller et al. (2019b) to the ‘pooled meiosis’ in which the crossovers from the two actual meioses have been combined, we find

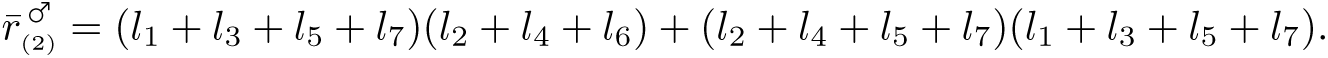

Across the sixteen possible segregation patterns (A*i*, B*j*), 𝔼[*IBD*_h-sib_] = 1/4 and

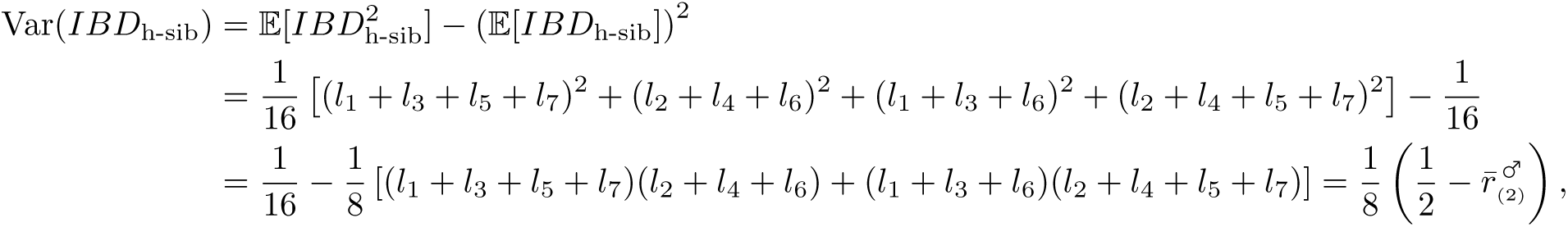

which is Eq. (3).

Like 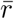, 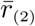 can be estimated from various kinds of data, including cytological data of crossover positions at meiosis I and sequence data from gametes. Using cytological data for human male spermatocytes from Lian et al. (2008), we construct a pooled meiosis from every possible pair of spermatocytes. Calculating the value of 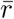 for each of these pooled meioses and averaging, we obtain 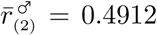 (MATLAB code for this calculation is available at github.com/cveller/IBD). Thus, from Eq. (3), the genetic relatedness of half-sibs who share a father but have unrelated mothers has variance 1.1 × 10^−3^ (*σ* = 0.033, CV ≈ 13%).

## 4 Application: Ancestry variance among F2s

A common experimental design involves mating individuals from two lines, populations, or species (A and B) to form a hybrid ‘F1’ generation, and then mating F1s to produce an F2 generation. Every F1 carries exactly one-half of its DNA from each species, but there is ancestry variance among F2s because of recombination and segregation in the F1s’ meioses (Hill 1993a).

Each F2 derives from an F1 mother’s egg and an F1 father’s sperm. Let the random variables 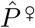 and 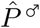 be the respective proportions of species-A DNA in the egg and sperm, and let *P* be the proportion of species-A DNA in an F2’s genome. Then 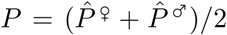, and, from Eq. (1), 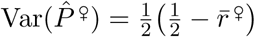 and 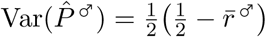. Finally, because 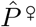 and 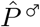 are independent, the ancestry variance among F2s is

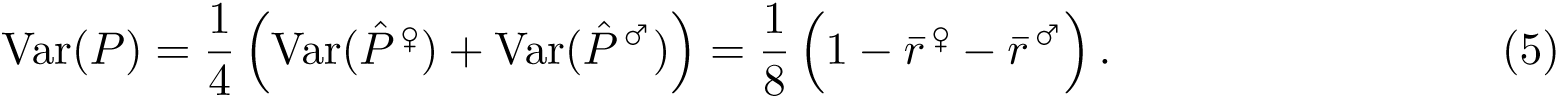

[If the F2s instead derived from a backcross of F1s to one of the parental species, the ancestry variance among F2s would be 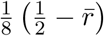, with 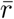 calculated for the sex of the F1s involved. This variance, and those among later generations of the backcrossing program, have been calculated by Hill (1993a) under the assumptions of a uniform recombination rate and no crossover interference.]

### Selection for one ancestry over the other

The calculation above assumes that no other forces are affecting ancestry among F2s. Such forces could include systematic selection among F2s in favor of alleles from one of the species, or meiotic drive in F1s, both of which would shift the distribution of ancestry among F2s towards one of the two species. For example, Matute et al. (2020) generated two crosses, each between a widely distributed species of *Drosophila* and a closely related island endemic, and observed in the resulting admixed populations that island ancestry was replicably selected against over time. If viability selection plays a role in this effect, then an ancestry skew towards the widespread species would be expected among adult F2s of these crosses.

An F2-based test for selection of this kind would involve comparing the observed average ancestry among F2s against the neutral null expectation of 1/2. In this case, Eq. (5) gives the appropriate null variance for the purpose of statistical inference; the standard error of the test is

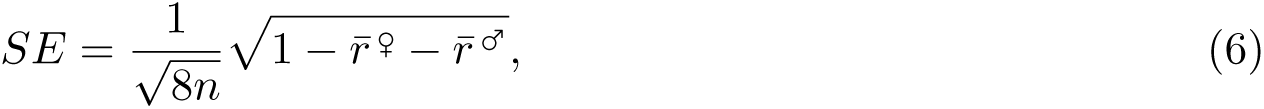

where *n* is the sample size of F2s for which ancestry proportions have been measured.

Substituting known values of 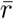 into Eq. (6) then shows us, for a given sample of F2s, how much their average ancestry proportion must deviate from 1/2 for us to reject the null hypothesis of neutrality. For example, using a linkage map generated from a cross of two closely related cichlid fish species (Feulner et al. 2018), we calculate a sex-averaged value of 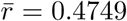. If ancestry fractions were measured for 10 F2s from this cross, then Eq. (6) tells us that a 4.9% or greater deviation of the average ancestry from the null expectation of 50% would be statistically distinguishable at the 5% significance level; if 100 F2s were assayed, the threshold detectable deviation would be 1.6%. Threshold deviations for a range of sample sizes are shown in Fig. 3 for the recombination processes of cichlids, humans, and *Drosophila melanogaster*. It is clearly seen that, because of *D. melanogaster ’*s low sex-averaged value of 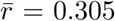 (see Discussion), much greater ancestry deviations among F2s are required for the null hypothesis of neutrality to be rejected, compared to cichlids and humans.

**Figure 3:**
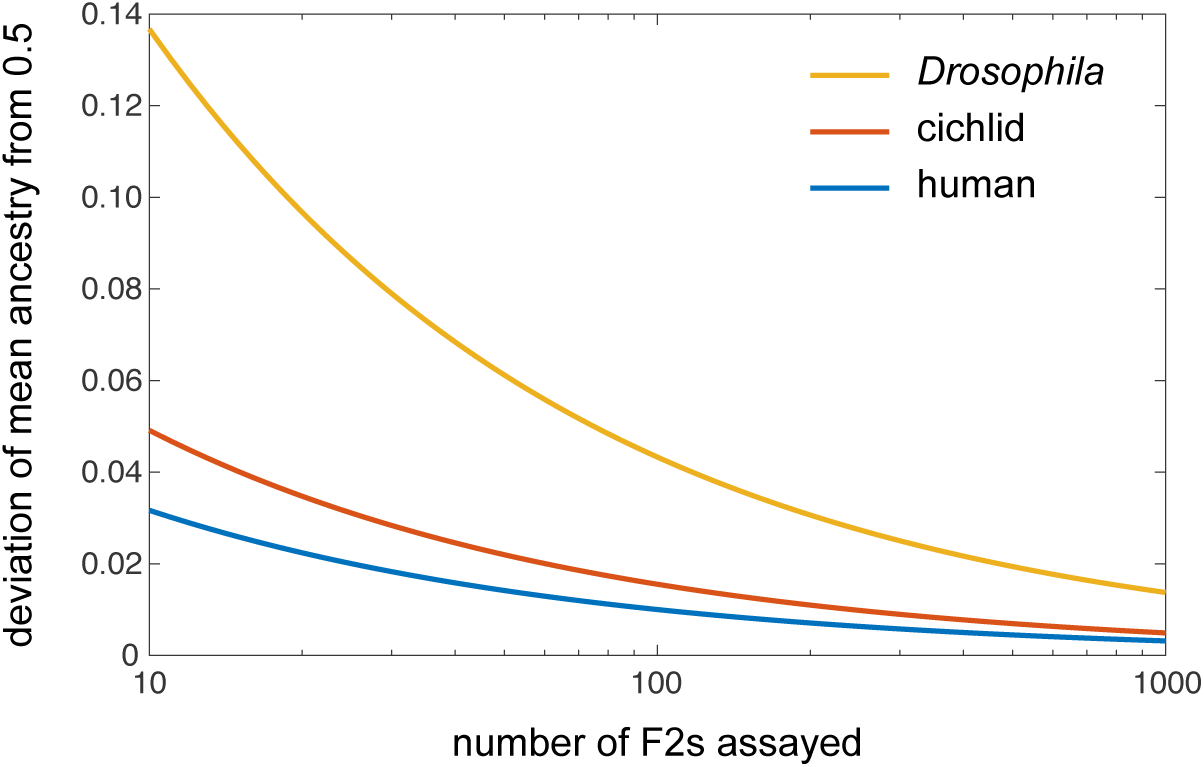
The minimum deviation of mean ancestry among F2s that can be statistically distinguished from the null expectation of 1/2 at the 5% significance level.

Note that the test described above can also be carried out for a specific genomic region of interest, by using region-specific values of 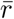. Alternatively, in a genome-wide scan, regions of the genome where ancestry deviations are particularly large can be statistically identified using region-specific values of 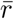 and correction for multiple hypothesis testing.

### Hybrid vigor

A common observation in hybrid crosses is heterosis, or hybrid vigor, whereby hybrids (especially F1s) are fitter than their parent lines. For example, if recessive deleterious mutations became fixed at different sites in species A and B, then F1s would be heterozygous at all of these sites, and would therefore enjoy higher fitness than their parents. Subsequently, among F2s, fitness depends on the number of these sites at which an individual is heterozygous for species ancestry.

To see what Eq. (5) can say about the operation of heterotic selection, let *π* be the probability that a randomly selected species-A allele differs from a randomly selected species-B allele at the same site—*π* is the average heterozygosity of F1s. Let the random variable *Ĥ* be the proportion of loci in an F2 zygote’s genome at which it carries alleles from both species, and let the random variable *H* be the proportion of loci heterozygous in an F2 zygote: *H* = π*Ĥ*. If there is no selection acting between the zygote stage and the stage at which F2s are genotyped, then the average proportion of loci that are heterozygous in an F2 at the time of genotyping is 𝔼[*H*] = *π*𝔼[*Ĥ*] = *π*/2, and, using a calculation similar to Eq. (3), the variance is

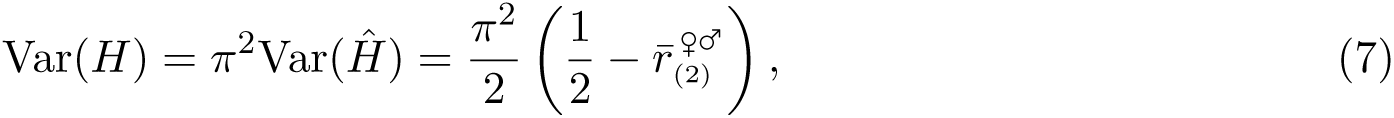

where 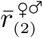 is the value of 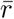 that results from pooling the crossovers of a random female meiosis and a random male meiosis. [Previous calculations of Var(*H*) have assumed that crossovers are uniformly distributed along chromosomes and do not exhibit interference—e.g., Franklin (1977, p. 67), where it is expressed as heterozygosity variance in the offspring of a selfing parent. More general quantities are calculated under the same assumptions by Franklin (1977); Weir et al. (1980); Hill and Weir (2011).]

Now let the random variable *H′* be the proportion of heterozygous loci in an F2 adult at the time of genotyping. If selection acts additively across loci so that an individual with a proportion *x* of heterozygous loci has relative viability 1 + *Sx*, then the variance in viability among F2 zygotes that is contributed by heterotic selection is *S*^2^Var(*H*), and it can be shown (SI Section S3) that the expected change in heterozygosity between the zygote and adult stages is

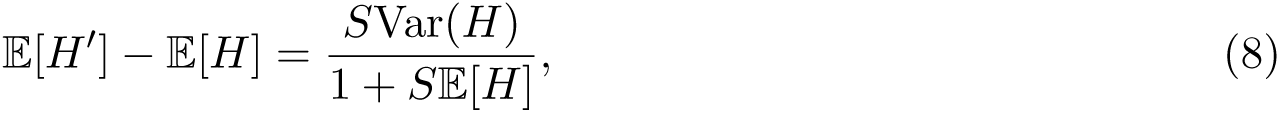

so that

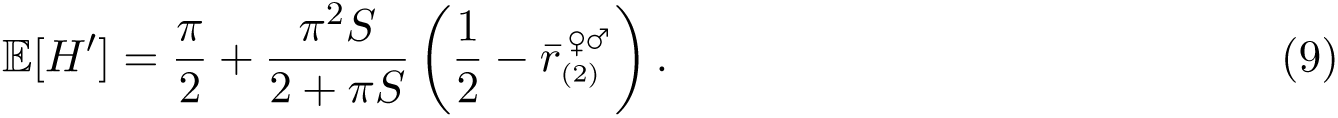

From this expression, an F2-based estimator for *S*, the strength of hybrid vigor, can be derived:

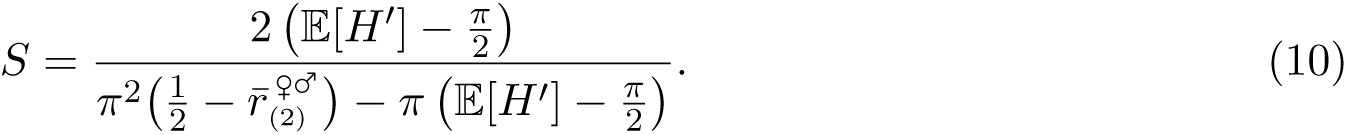

### Bateson-Dobzhansky-Muller incompatibilities

A further form of selection that can act among F2s is epistasis between ‘incompatible’ pairs of alleles from the two species, acting at distinct loci. The effect of these ‘Bateson-Dobzhansky-Muller incompatibilities’ (BDMIs) on ancestry among adult F2s will depend in a complicated way on patterns of (cross-locus) dominance of the alleles involved, as well as recombination in F1s (Turelli and Orr 2000). Nevertheless, it turns out that the strength of selection against BDMIs can be characterized in terms of aggregate recombination, and specifically the metrics 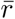 and 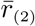. Owing to the complexity of this case (including an important role for sex chromosomes), we shall treat it in more detail elsewhere.

## 5 Discussion

Relatives of a given pedigree relationship vary in how much of their DNA they share identically by descent, because of variable patterns of recombination and segregation in their pedigrees. We have shown that the variance of genetic relatedness is determined by aggregate recombination, as quantified by 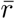 and analogous metrics (i.e., averages of functions of pairwise recombination rates). Therefore, features of the aggregate recombination process that affect these metrics—such as the number of chromosomes and heterogeneity in their size, the number of crossovers and their location along chromosomes, and the spatial organization of crossovers with respect to one another—also affect variance in genetic relatedness. Below, we discuss how the influence of these meiotic features on variance in genetic relatedness can be understood, perhaps more intuitively, by thinking in terms of their effect on aggregate genetic shuffling.

### Sex differences in recombination

In many species, male and female meiosis differ both in the number and location of crossovers [reviewed by Lenormand and Dutheil (2005); Sardell and Kirkpatrick (2020)]. In human male meiosis, crossovers are fewer and more terminally localized than in female meiosis. Both factors decrease the total amount of genetic shuffling (Veller et al. 2019b). This explains the observation of Caballero et al. (2019) that relatives who are related predominantly via males have a higher variance of genetic relatedness than relatives related predominantly via females.

Such effects will be especially pronounced in species where one sex has no crossing over in meiosis. For example, in *Drosophila*, there is crossing over in oogenesis but not in spermatogenesis. Using chromosome lengths from Release 6 of the *D. melanogaster* reference genome (Hoskins et al. 2015) and the female linkage map produced by Comeron et al. (2012), we calculate autosomal values of 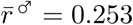 and 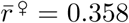 (the sex-averaged value is 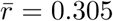). Substituting these values into Eq. (2), we find that the variances of (autosomal) relatedness to a paternal and a maternal grandparent are 0.0308 and 0.0156, respectively (*σ* = 0.175 and 0.125).

### Crossover interference

It has recently been shown, by computer simulation of various forms of crossover patterning along chromosomes, that crossover interference tends to decrease the variance of genetic relatedness between relatives (Caballero et al. 2019). Veller et al. (2019b) demonstrated that interference among crossovers increases the amount of genetic shuffling that they cause (increasing 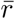 and analogs). The intuition is that, when two crossovers occur very close to each other, they cancel each other’s effect on genetic shuffling, behaving more like a single crossover. Such ‘stepping on toes’ is prevented by crossover interference, thus increasing genetic shuffling. This provides an intuitive explanation of the result of Caballero et al. (2019).

For example, using a simulation method employed by Mancera et al. (2008) and Wang et al. (2012) to resample empirically observed crossovers in an interference-less way, Veller et al. (2019b) calculated that, in human male, interference among crossovers increases their contribution to 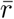 by about 15%. By this measure, crossover interference in human male meiosis decreases the variance of genetic relatedness to a paternal grandparent from 1.82 × 10^−3^ to 1.59 × 10^−3^ [Eq. (2)], a decrease of 12.6%.

### Recombination hotspots

White and Hill (2020) studied the effect of adding a recombination hotspot to a chromosome on the variance of genetic relatedness between relatives, and concluded that it depends on the position of the hotspot along the chromosome. This can be understood in terms of the effect of crossover position on 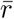—terminal crossovers cause little genetic shuffling. I.e., this result of White and Hill (2020) is an observation about the broad-scale distribution of crossovers.

A separate question about hotspots concerns their effect on genetic relatedness if the broad-scale distribution of crossovers is held constant. Thinking in terms of aggregate genetic shuffling suggests that the effect of hotspots in this case will depend on the pedigree relationship in question. For example, in the presence of crossover interference, hotspots will have little effect on 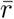, since 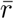 is determined by the broad-scale distribution of crossovers in individual meioses; thus, hotspots will have little effect on genetic relatedness between grandparent and grandoffspring [Eq. (2)]. However, hotspots are expected to decrease the value of 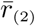 because, in the pooling of crossovers from two independent meioses, crossovers from the two meioses will sometimes land in the same hotspot, thus ‘wasting’ one of the crossovers. Therefore, confining crossovers to small regional hotspots is expected to increase the variance of genetic relatedness between siblings [Eq. (4)], and other pedigree relationships involving multiple meioses. The size of the effect will depend on the density and strength of the hotspots.

### Crossover covariation

It has recently been shown across diverse eukaryotes that the number of crossovers per chromosome covaries positively across chromosomes within individual meiotic nuclei (Wang et al. 2019). This ‘crossover covariation’ substantially increases the variance of crossover number per gamete, which clearly affects the distribution of genetic relatedness among relatives. However, because crossover covariation does not change the probability that a given pair of loci are recombinant in a gamete, it does not alter 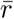 or analogs (since these are averages of functions of pairwise recombination rates—see SI Sections S1 and S2). Therefore, crossover covariation affects neither the mean nor the variance of genetic relatedness among relatives, but it will affect higher-order moments.

## 6 Conclusion

The quantities considered in this paper—variances of genetic relatedness and of ancestry in hybrid crosses—play an important role in many applications in genetics (Thompson 2013). For this reason, there exists a substantial prior literature calculating these quantities [e.g., Franklin (1977); Hill (1993a); Hill and Weir (2011)]. In this literature, it is generally assumed that crossovers are uniformly distributed along chromosomes, and that crossover interference is absent [though see White and Hill (2020); Caballero et al. (2019)]. These assumptions allow the variance of genetic relatedness to be expressed as a function of the total map lengths of individual chromosomes (Hill and Weir 2011), which is useful when these are the only quantities that can be reliably estimated given available data (e.g., low-density linkage maps). Nevertheless, it has been shown that violation of the assumptions of a uniform recombination rate and no crossover interference, as will generally be the case, can lead to substantially biased estimates of the variance of genetic relatedness (Caballero et al. 2019; White and Hill 2020).

We have shown that, when no assumptions are made about the aggregate recombination process, the variance of genetic relatedness is the average of some function of pairwise recombination rates. For the particular pedigree relationships of grandparent-grandoffspring and siblings, the variance of genetic relatedness is a function of 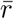, the average proportion of locus pairs that recombine in meiosis. Since these quantities—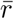 and analogs—can readily be estimated from modern cytological and genetic data (Veller et al. 2019b), it will now be possible in many cases to calculate the variance of genetic relatedness in a fully unbiased way.

## Acknowledgements

We are grateful to Nicolas Bierne, Erin Calfee, Graham Coop, Jim Mallet, Molly Schumer, and Sivan Yair for helpful discussions and comments.

## S1 General case for direct descent

Label the starting generation 0, so that the offspring generation is 1, the grand-offspring generation 2, etc. For an individual in generation 0, one of its descendants in generation *t*, and a locus *k*, let 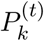 be a random variable that takes the value 1 if an allele carried by the generation-*t* descendant at locus *k* was inherited from the generation-0 individual, and takes the value 0 otherwise. Clearly,

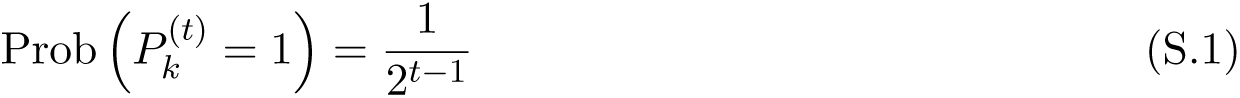

and because 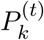 can take on only the values 0 or 1, 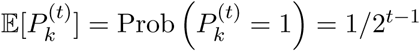. Then

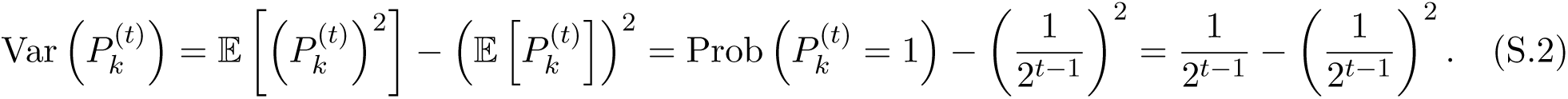

### Case 1: No sex differences in recombination

Consider two loci, *i* and *j*. For the alleles at these loci in the generation-*t* descendant both to have been inherited from the generation-0 ancestor requires that the loci never have been recombinant in any of the gametes linking the generation-*t* descendant and the specified generation-0 ancestor (probability 1 − *r*_*ij*_ for each relevant gamete, starting with that produced by the generation-1 descendant and ending with that produced by the generation-[*t* − 1] descendant) and, conditional on this, that the appropriate alleles co-segregated to the gamete in each relevant meiosis (probability 1/2 each time). Therefore,

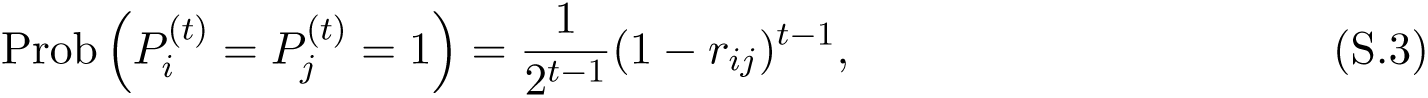

so that

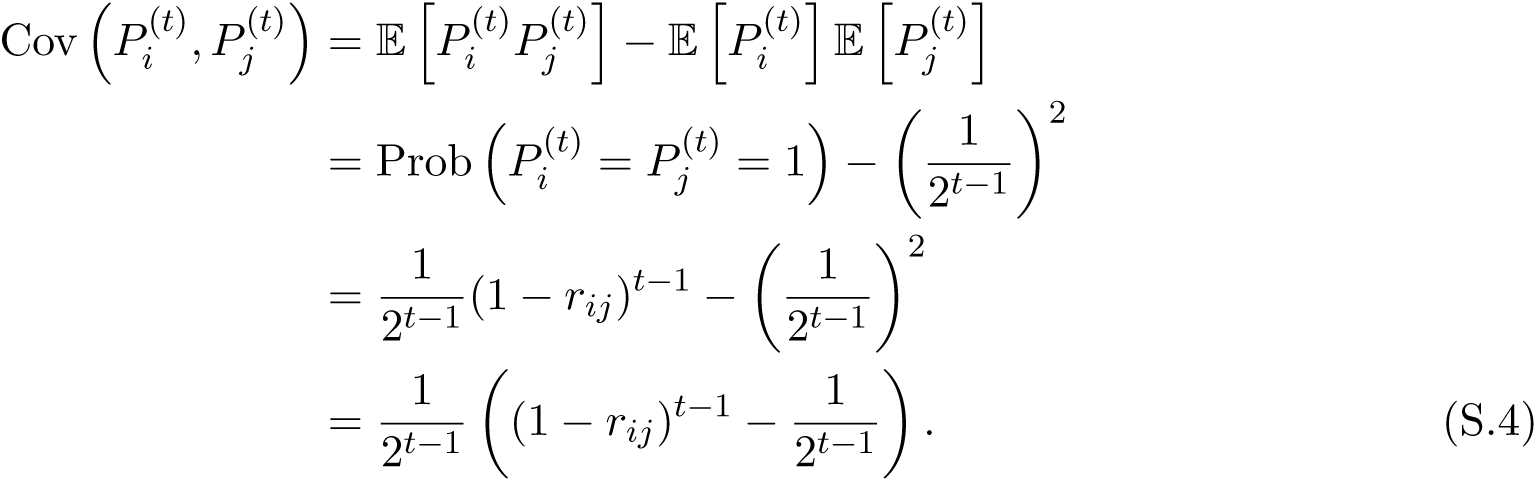

Assume that there are *L* loci in total, with *L* very large, and let *P* ^(*t*)^ be the proportion of the the generation-*t* descendant’s genome inherited from the generation-0 ancestor: 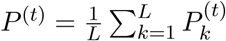. Then

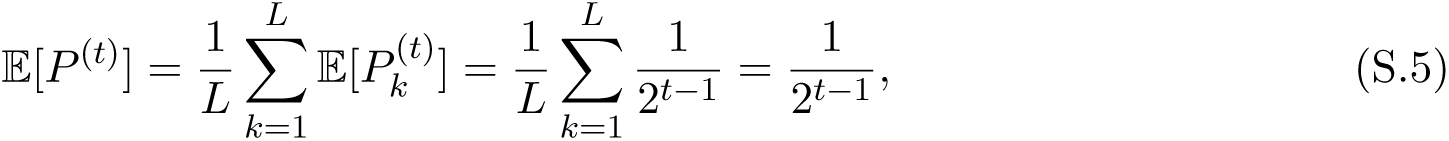

and

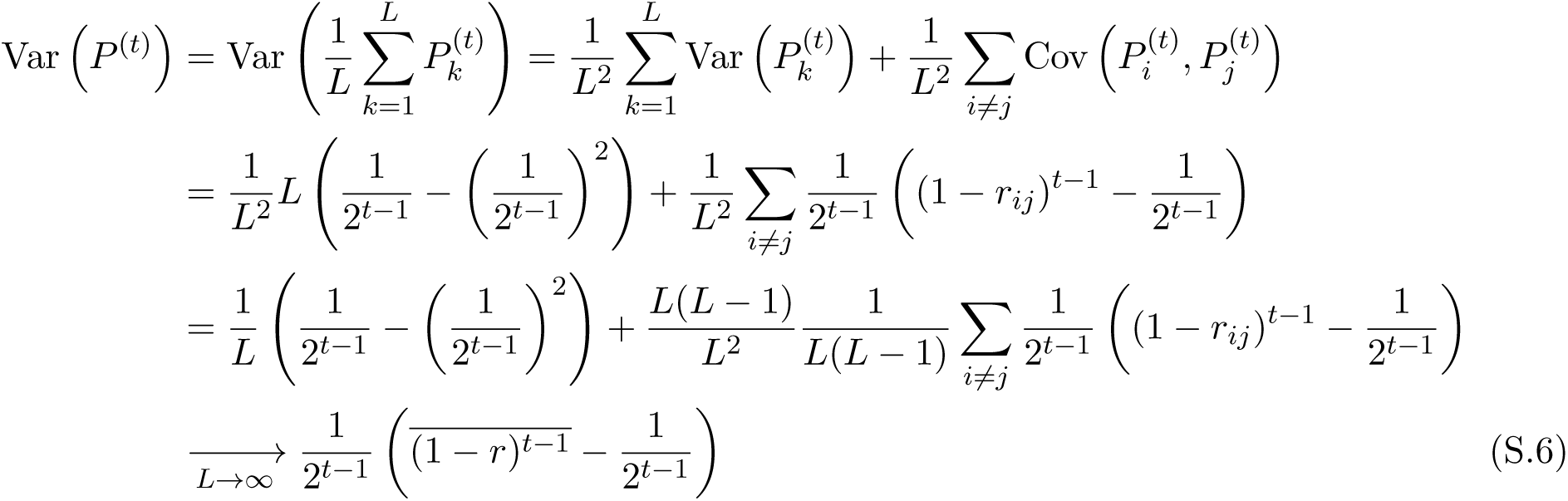

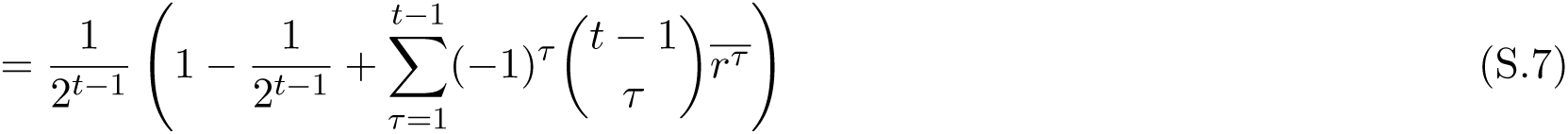

where a bar represents the average taken with respect to all locus pairs, and 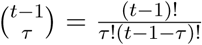. The limit follows from the fact that 1/*L* → 0, *L*(*L* − 1)/*L*^2^ → 1, and the number of pairs (*i, j*) such that *i* ≠ *j* is *L*(*L* − 1).

Finally, let *IBD*^(*t*)^ be the fraction of the generation-*t* individual’s (diploid) genome that is inherited identically by descent from the generation-0 ancestor. Clearly *IBD*^(*t*)^ = *P* ^(*t*)^/2, and so

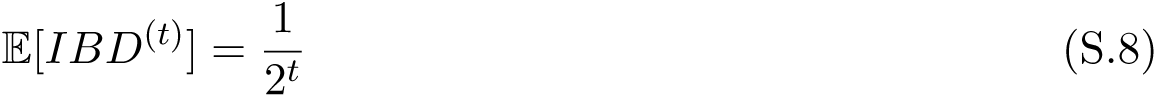

and

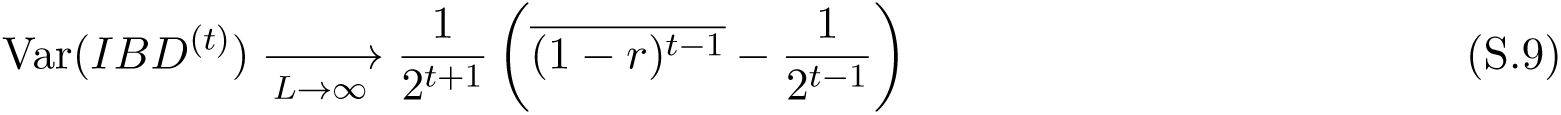

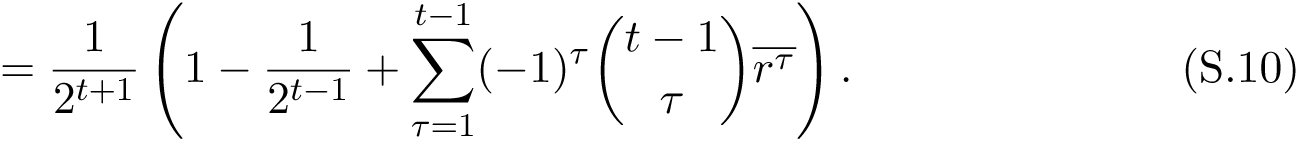

In the special case of the descendant being a grand-offspring (*t* = 2), Eq. (S.9) becomes

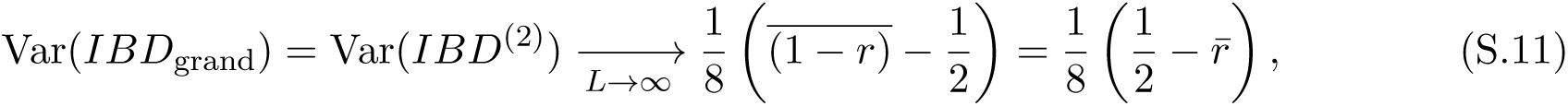

which is Eq. (2) in the Main Text.

### Case 2: Sex differences in recombination

Let 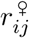 and 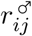 be the sex-specific recombination rates between loci *i* and *j*. If, among the *t* − 1 individuals in the lineage between the generation-0 ancestor and the focal generation-*t* descendant, there are *f* females and *m* = *t* − 1 − *f* males, then

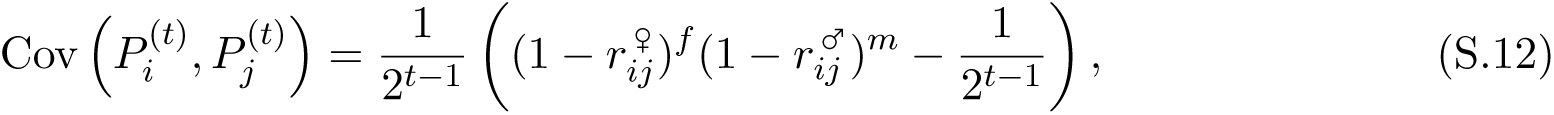

so that, by a similar calculation to Eq. (S.9) above,

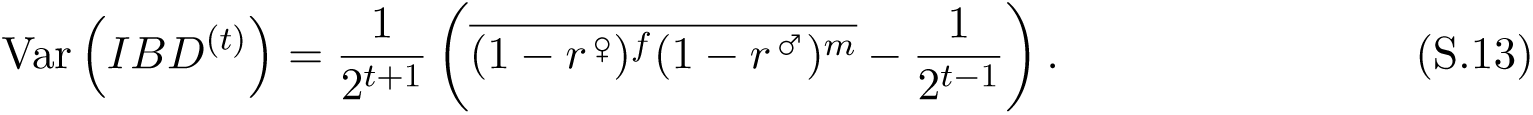

If the number of females in the lineage is not known, it can be taken to be binomially distributed with parameter 1/2, in which case the average in Eq. (S.13) is calculated across all locus pairs and all possible numbers of females *f* = 0, 1, …, *t* − 1 (with associated probabilities 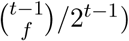).

## S2 General case for indirect relationships

### Relationships with one common ancestor

Consider an individual (generation 0) and two of its descendants (generation *t*_1_ and *t*_2_) who have no more recent common ancestor than the generation-0 individual, and no other recent common ancestor that is not also an ancestor of their shared generation-0 ancestor [FIGURE]. The two generation-1 ancestors of the focal descendants (which could be the focal descendants themselves if *t*_1_ = *t*_2_ = 1) are half-sibs. Let 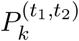 be a random variable that takes on the value 1 if both focal descendants carry, at locus *k*, an allele inherited identically from their common generation-0 ancestor. Assuming Mendelian segregation,

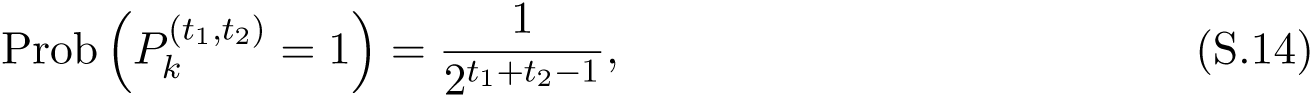

so that 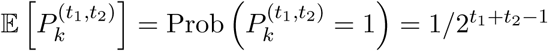 and

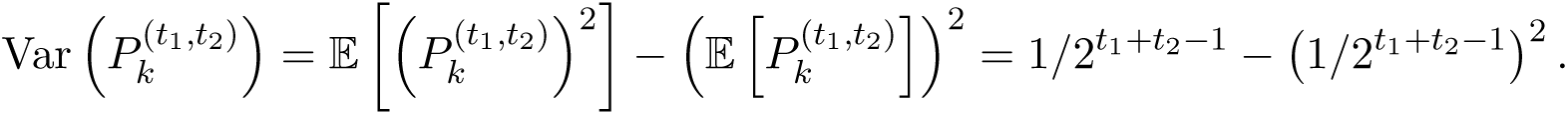

Now consider two loci, *i* and *j*. For the alleles at both loci in both descendants to have been inherited from their common ancestor in generation 0 (i.e., for the individuals to be IBD at these two loci) requires (i) that the two generation-1 ancestors be IBD at the two loci, which, because they are half-sibs, occurs with probability 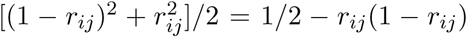, (ii) that the two loci not be recombinant in any subsequent gamete leading to the focal generation-*t*_1_ and generation-*t*_2_ descendants, which occurs with probability 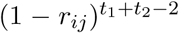, and (iii) that, given (i) and (ii), the ancestor’s alleles always segregate into the gametes leading to the focal descendants, which occurs with probability 1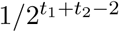. Therefore,

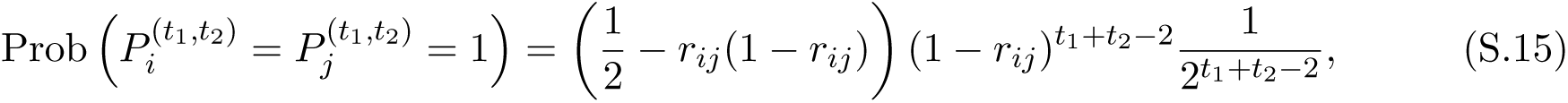

so that

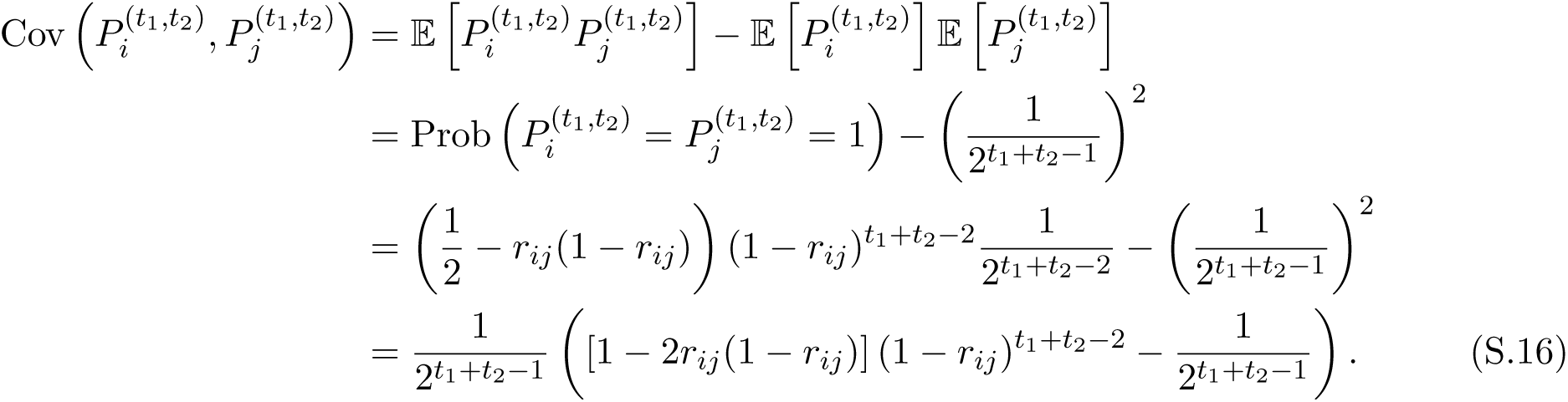

Now let 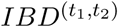 be the fraction of the genome that both the focal descendants have inherited from their common generation-0 ancestor: 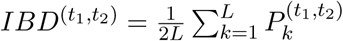. Then

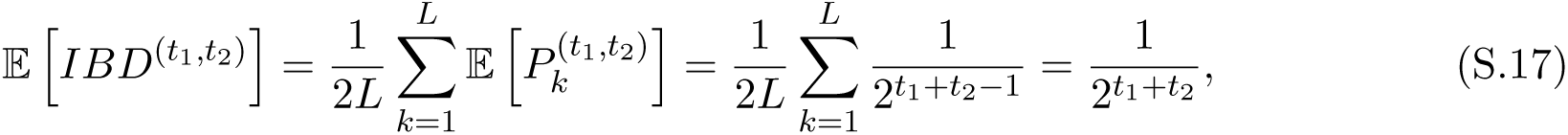

while

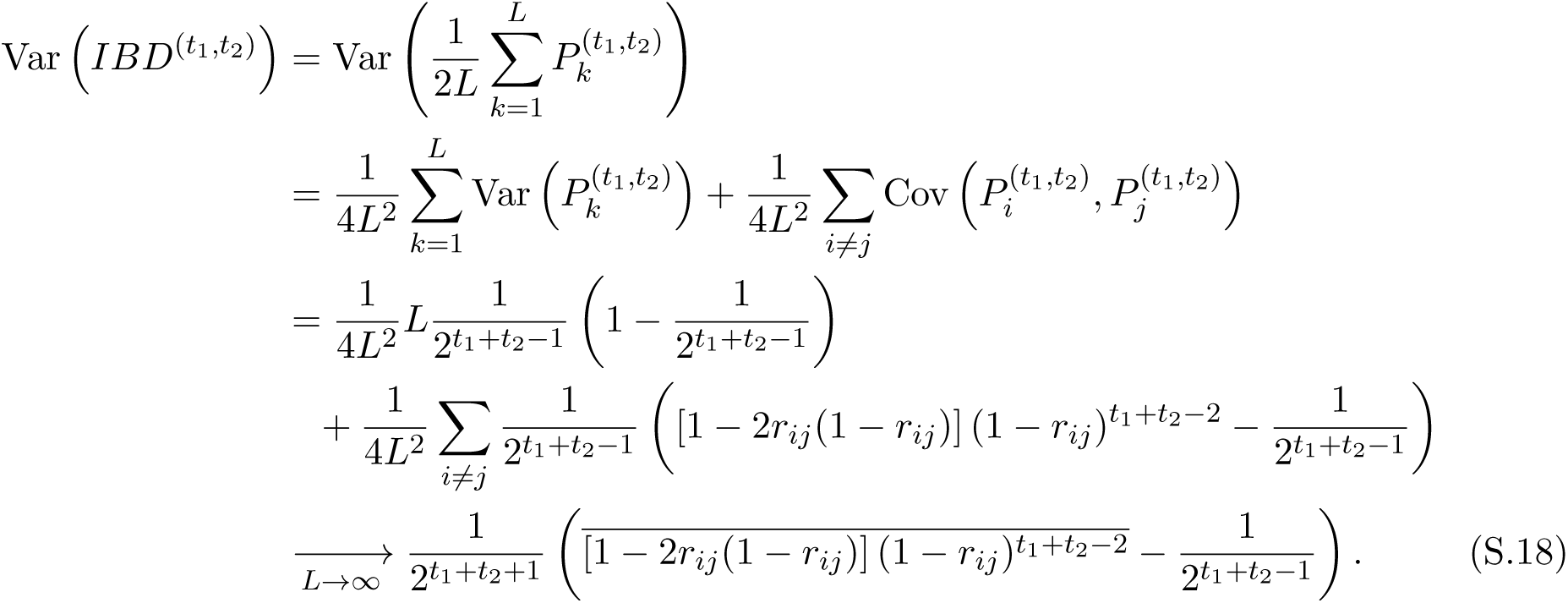

In the special case of the focal descendants being half-sibs (*t*_1_ = *t*_2_ = 1), Eq. (S.18) becomes

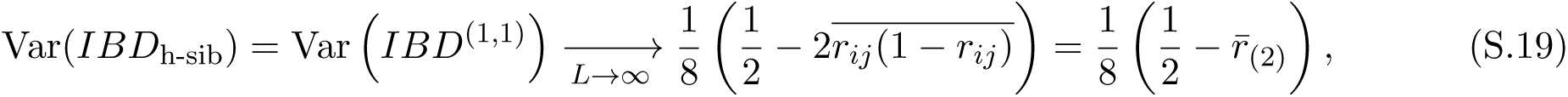

which is Eq. (3) in the Main Text. Here, 2*r*_*ij*_(1 − *r*_*ij*_) is the probability that *i* and *j* are recombinant in exactly one of two gametes, and 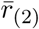 is the average value of 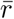 calculated from the pooled crossovers of two independent meioses of the common parent.

### Relationships with two common ancestors

Consider an unrelated mating pair (generation 0) and two of its descendants (generation *t*_1_ and *t*_2_) who have no more recent common ancestors than the generation-0 mating pair, and no other recent common ancestor that is not an ancestor of one member of the generation-0 mating pair. Note that, if *t*_1_ = *t*_2_ = 1, the focal descendants are full-siblings. Thus, we have restricted attention to two-ancestor pedigrees where the two ancestors were a mating pair; i.e., we have excluded from attention pedigrees such as that depicted in Fig. S1B below.

Let 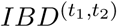 be the proportion of the focal descendants’ genomes that they share IBD. Label the members of the mating pair 1 and 2 (female and male, respectively), and let 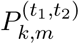 be a random variable that takes the value 1 if both focal descendants carry, at locus *k*, an allele inherited from member *m* ∈ (0, 1) of the mating pair. Then 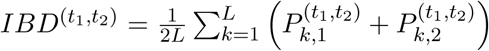. Notice that, if *t*_1_, *t*_2_ 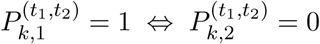. Therefore, we consider three separate cases:

**Figure S1:**
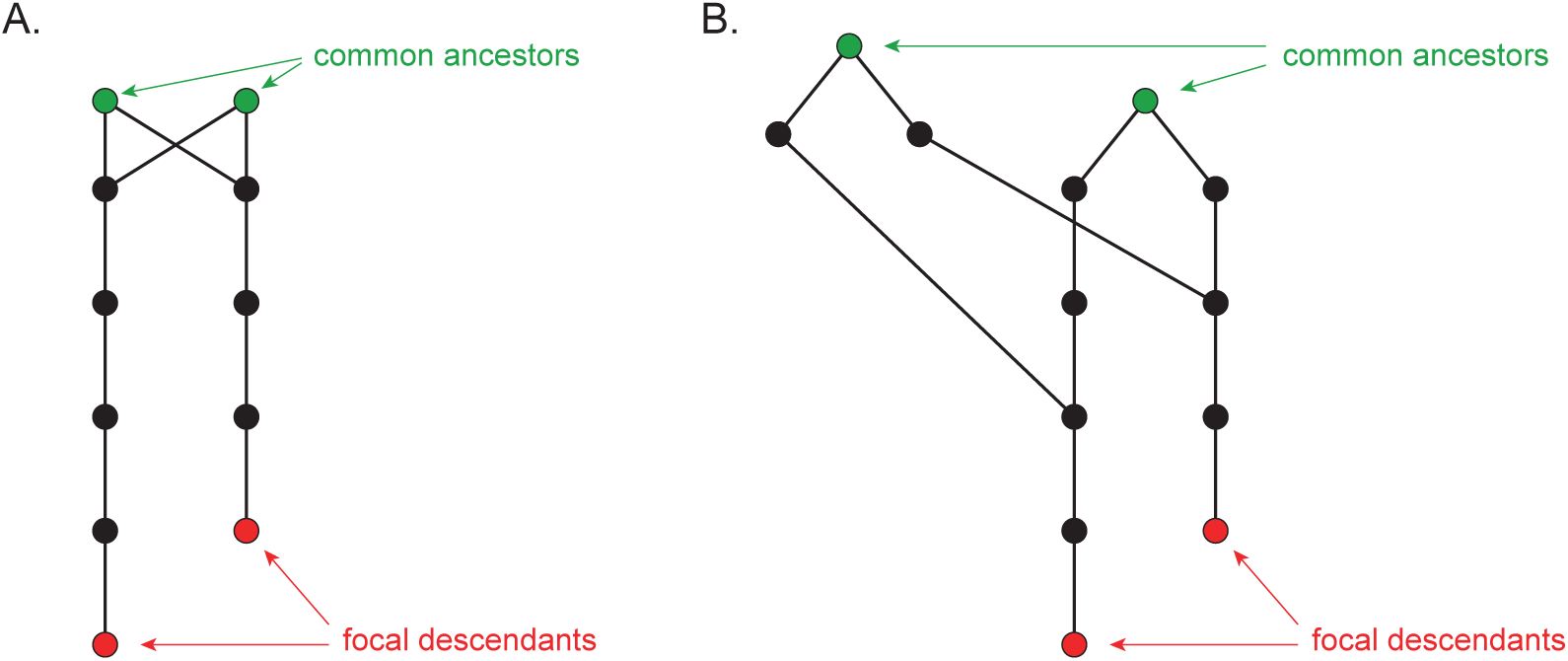
Examples of the general kinds of two-ancestor pedigrees we do consider (A) and do not consider (B). In pedigree A, *t*_1_ = 5 and *t*_2_ = 4.

### Case 1: *t*_1_ = *t*_2_ = 1 (full-sibs)

In the case of full-sibs, because they each inherit their maternal and paternal genomes independently, the random variables 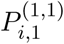 and 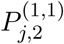 are independent for all *i, j*, and have the same distribution as 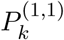 defined in the subsection above (Relationships with one common ancestor). Moreover, the random variables 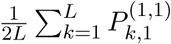 and 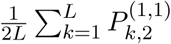 have the same distribution as 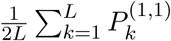, with the appropriate recombination process—female and male, respectively—substituted in each case. Therefore,

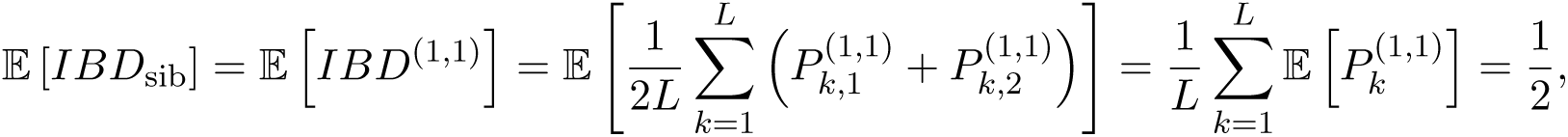

and

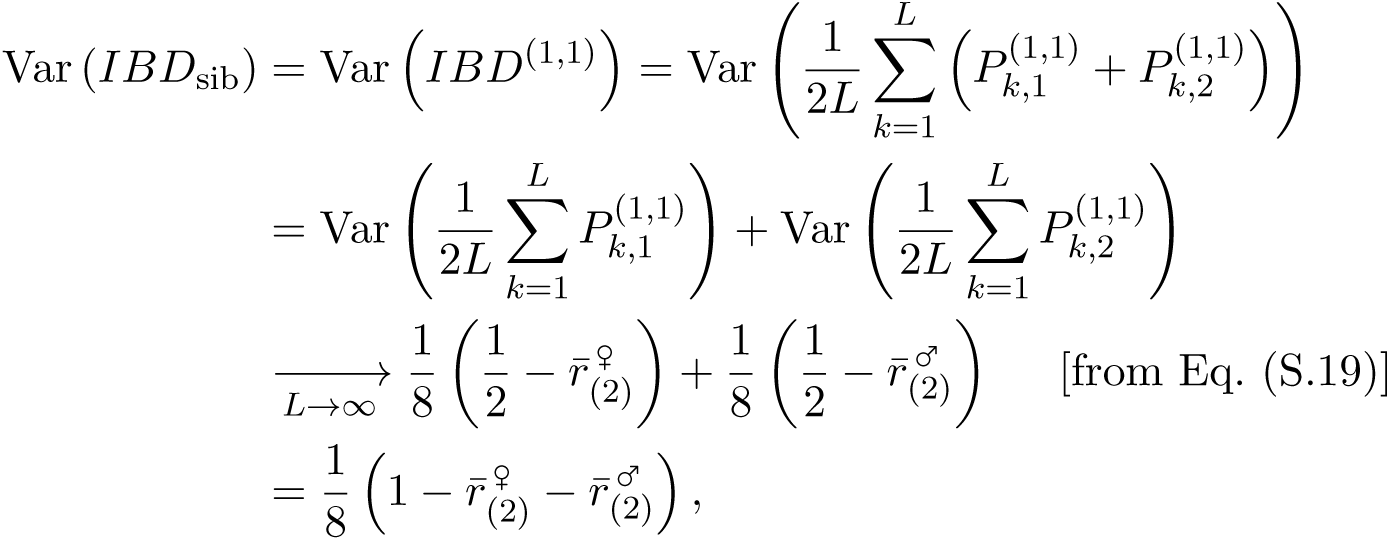

which is Eq. (4) in the Main Text.

### Case 2: *t*_1_ = 1, *t*_2_ > 1

Without loss of generality, let the first focal individual be an offspring of the ancestral mating pair, and the second focal individual a more distant descendant. Then the second individual carries one haploid copy of the genome inherited through the pedigree, while the first individual carries two copies. Let the random variable 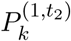 take on the value 1 if the focal descendants carry, at locus *k*, an allele inherited IBD. Clearly 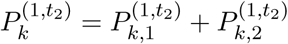, so that

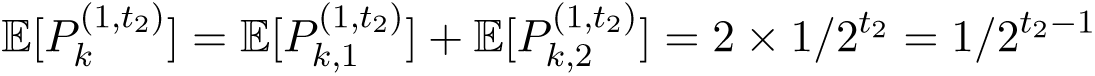

and 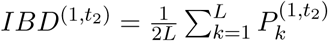. Now consider two loci, *i* and *j*. For 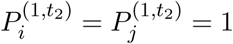, we require one of the following mutually exclusive events to occur:

- 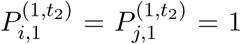. This requires that the generation-1 siblings inherited the same maternal alleles at *i* and *j* (probability 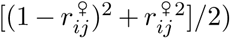, and that this allele pair was thereafter transmitted faithfully to the focal generation-*t*_2_ descendant (probability 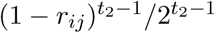. **Total probability:** 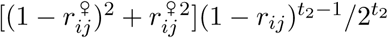.
- 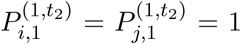. This requires that the generation-1 siblings inherited the same paternal alleles at *i* and *j* (probability 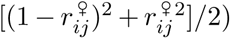), and that this allele pair was thereafter transmitted faithfully to the focal generation-*t*_2_ descendant (probability 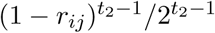. **Total probability:** 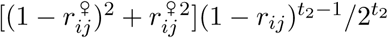.
- 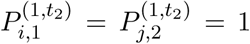. This requires that the generation-1 siblings inherited the same maternal allele at *i* (probability 1/2) and the same paternal allele at *j* (probability 1/2), that these two alleles were transmitted together to the generation-2 individual in the lineage of the focal generation-*t*_2_ descendant (probability *r*_*ij*_/2), and that these two alleles were then transmitted together to the focal generation-*t*_2_ descendant (probability 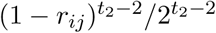. **Total probability:** 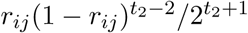.
- 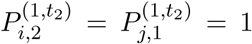. This requires that the generation-1 siblings inherited the same paternal allele at *i* (probability 1/2) and the same maternal allele at *j* (probability 1/2), that these two alleles were transmitted together to the generation-2 individual in the lineage of the focal generation-*t*_2_ descendant (probability *r*_*ij*_/2), and that these two alleles were then transmitted together to the focal generation-*t*_2_ descendant (probability 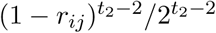. **Total probability:** 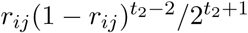.

Therefore,

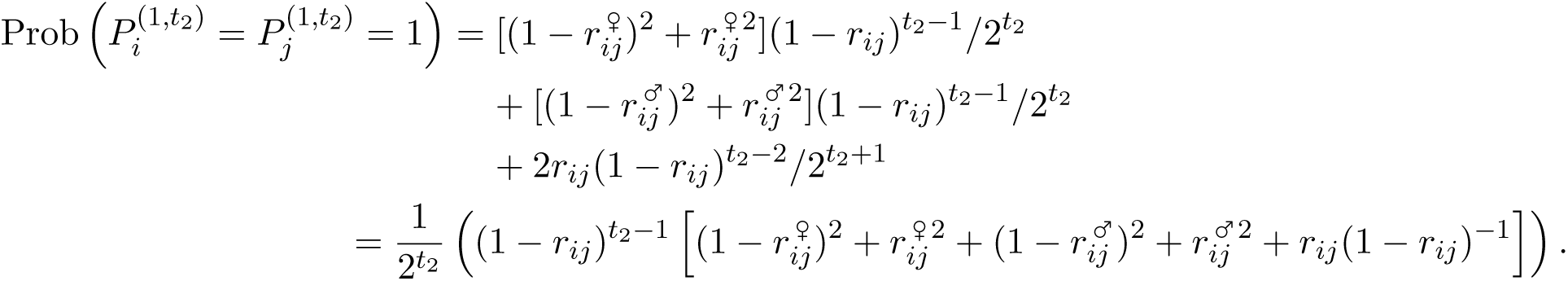

If the sex of members of the pedigree other than the ancestral mating pair is known, then it is clear where in the above calculations to substitute specific male or female values of *r*_*ij*_.

From the calculations above,

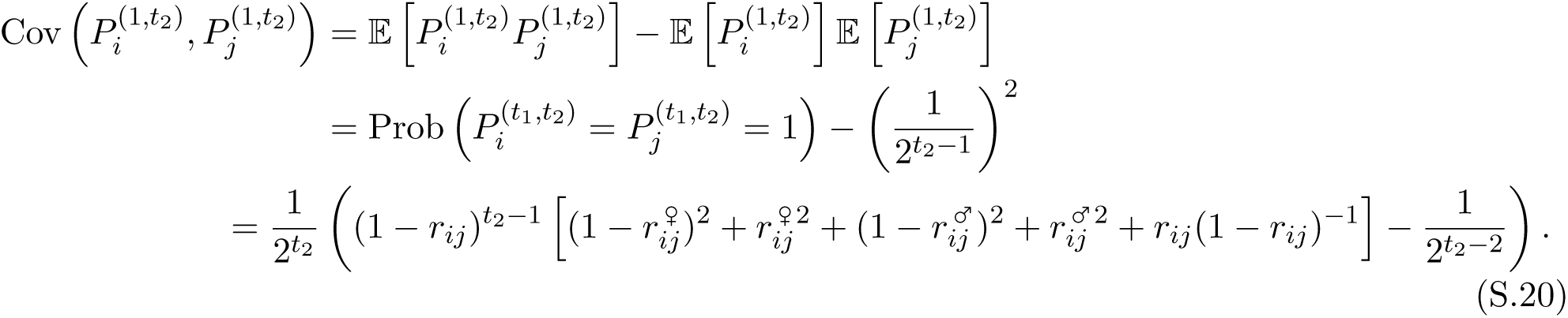

Using this result, and from calculations similar to those throughout this paper,

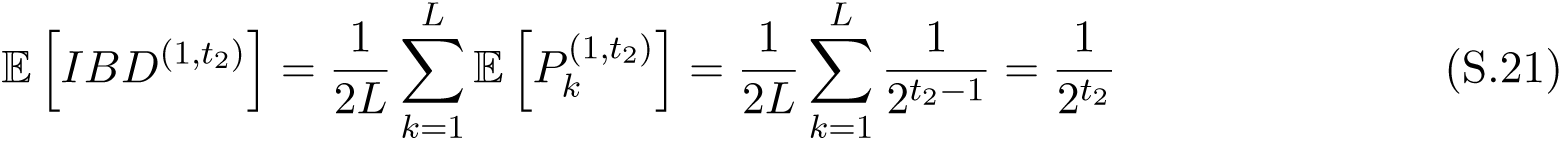

is the coefficient of relationship (e.g., 1/4 for aunt-nephew [*t*_2_ = 2]), while

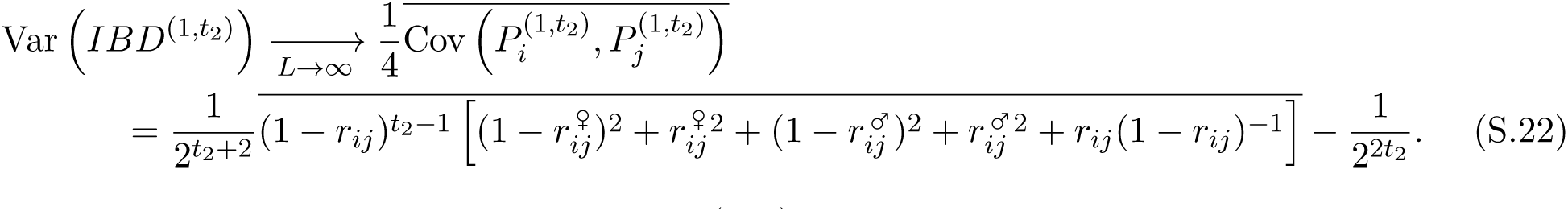

### Case 3: *t*_1_, *t*_2_ > 1

Let the random variable 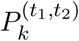 take on the value 1 if the focal descendants have inherited, within the focal pedigree, the same allele at locus *k*. Then

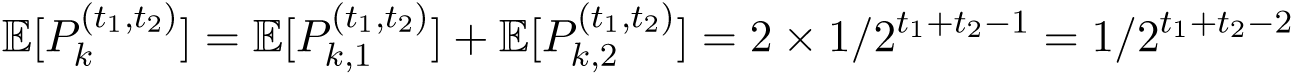

and 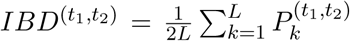. Now consider two loci, *i* and *j*. For 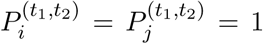, we require one of the following mutually exclusive events to occur:

- 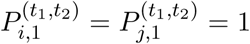. This requires that the generation-1 siblings inherited the same maternal alleles at *i* and *j* (probability 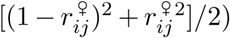, and that this allele pair was thereafter transmitted faithfully to both focal descendants (probability 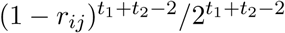. **Total probability:** 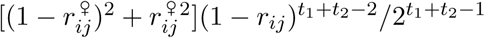.
- 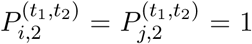. This requires that the generation-1 siblings inherited the same paternal alleles at *i* and *j* (probability 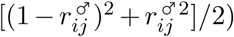, and that this allele pair was thereafter transmitted faithfully to both focal descendants (probability 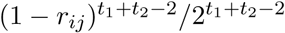. **Total probability:** 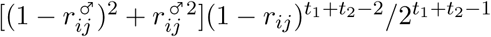.
- 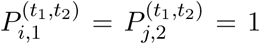. This requires that the generation-1 siblings inherited the same maternal allele at *i* (probability 1/2) and the same paternal allele at *j* (probability 1/2), that both generation-1 siblings transmitted these two alleles in producing the generation-2 cousins (probability *r*^2^/4), and that the allele pair was then transmitted faithfully to both focal descendants (probability 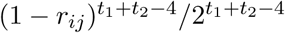. **Total probability:** 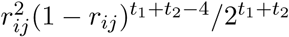.
- 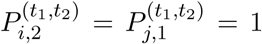. This requires that the generation-1 siblings inherited the same paternal allele at *i* (probability 1/2) and the same maternal allele at *j* (probability 1/2), that both generation-1 siblings transmitted these two alleles in producing the generation-2 cousins (probability 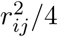), and that the allele pair was then transmitted faithfully to both focal descendants (probability 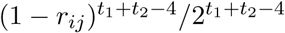. **Total probability:** 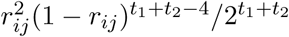.

Therefore,

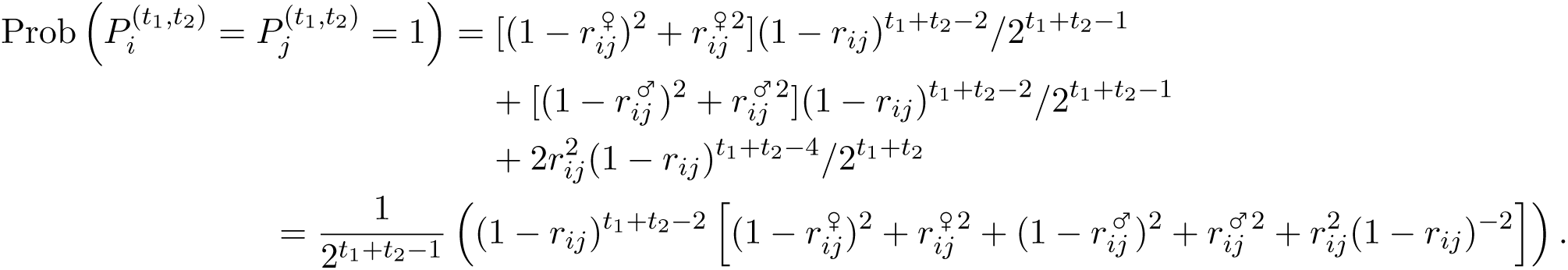

Again, if the sex of members of the pedigree other than the ancestral mating pair is known, then it is clear where in the above calculations to substitute specific male or female values of *r*_*ij*_.

From the calculations above,

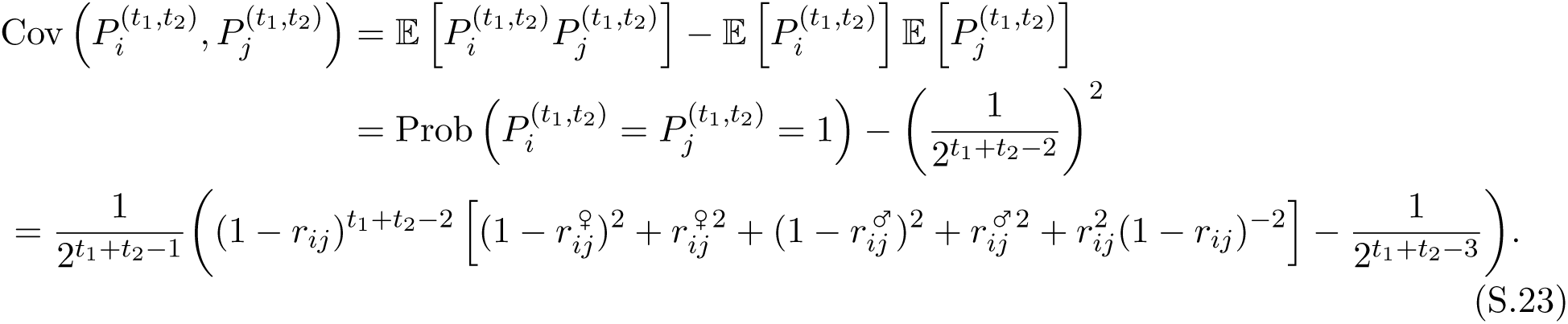

Using this result, and from calculations similar to those throughout this paper,

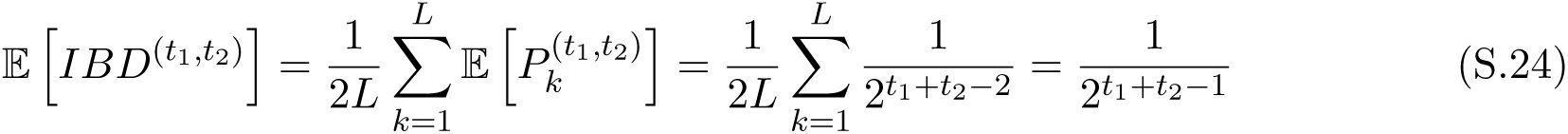

is the coefficient of relationship (e.g., 1/8 for full cousins [*t*_1_ = *t*_2_ = 2]), while

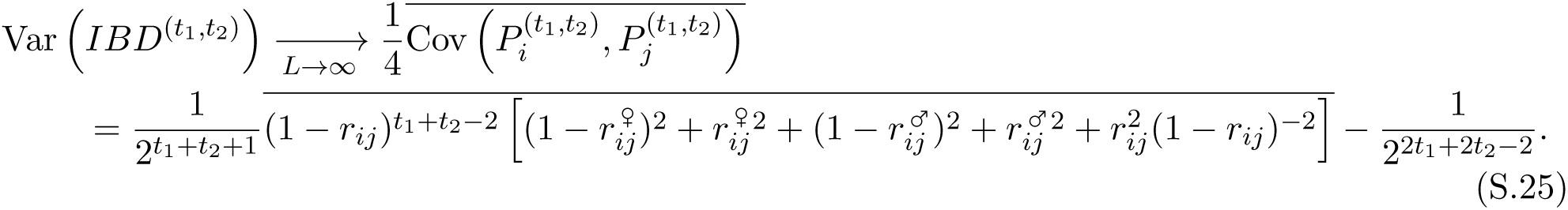

## S3 Relationship between variance in heterozygosity and change in heterozygosity under hybrid vigor

Let the random variable *H* be the proportion of loci that are heterozygous in a zygote. Suppose that selection acts according to a scheme of hybrid vigor, where an individual with proportion *x* of loci heterozygous has relative viability 1 + *Sx*, where viability here refers to the probability of surviving from zygote stage to the stage at which genotyping of F2s occurs. The random variable *H′* is the proportion of loci that are heterozygous at the stage of genotyping. Let the probability density functions for *H* and *H′* be *f* (*x*) and *g*(*x*) respectively. Then

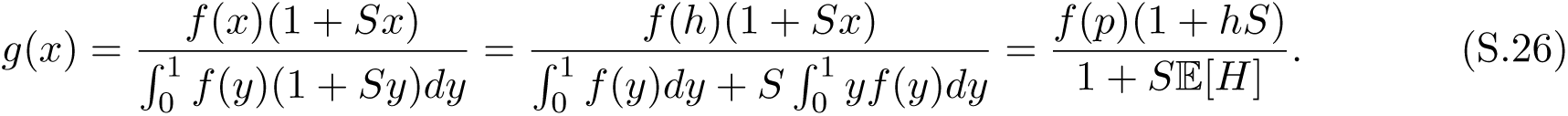

From this,

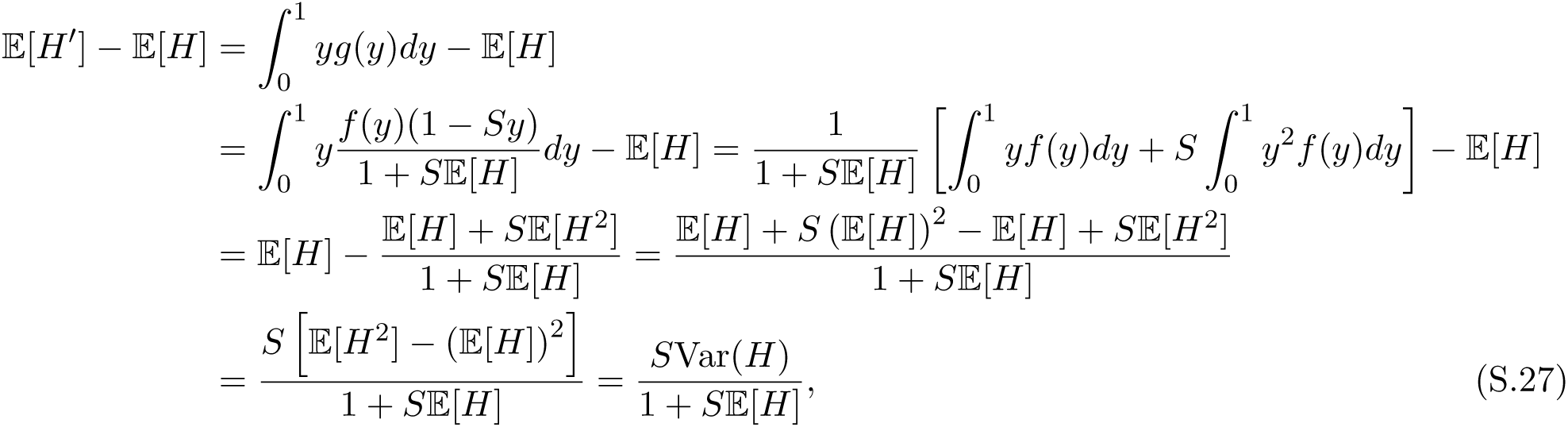

which is Eq. (8) in the Main Text.

## References

Caballero, M., Seidman, D. N., Qiao, Y., Sannerud, J., Dyer, T. D., Lehman, D. M., Curran, J. E., Duggirala, R., Blangero, J., Carmi, S., et al. (2019). Crossover interference and sex-specific genetic maps shape identical by descent sharing in close relatives. PLoS Genetics, 15(12):e1007979.

Comeron, J. M., Ratnappan, R., and Bailin, S. (2012). The many landscapes of recombination in *Drosophila melanogaster*. PLoS Genetics, 8(10):e1002905.

Feulner, P. G. D., Schwarzer, J., Haesler, M. P., Meier, J. I., and Seehausen, O. (2018). A dense linkage map of Lake Victoria cichlids improved the *Pundamilia* genome assembly and revealed a major QTL for sex-determination. G3: Genes, Genomes, Genetics, 8(7):2411–2420.

Franklin, I. R. (1977). The distribution of the proportion of the genome which is homozygous by descent in inbred individuals. Theoretical Population Biology, 11(1):60–80.

Gorlov, I. P. and Gorlova, O. Y. (2001). Cost–benefit analysis of recombination and its application for understanding of chiasma interference. Journal of Theoretical Biology, 213(1):1–8.

Guo, S.-W. (1996). Variation in genetic identity among relatives. Human Heredity, 46(2):61–70.

Hill, W. G. (1993a). Variation in genetic composition in backcrossing programs. Journal of Heredity, 84(3):212–213.

Hill, W. G. (1993b). Variation in genetic identity within kinships. Heredity, 71(6):652–653.

Hill, W. G. and Weir, B. S. (2011). Variation in actual relationship as a consequence of Mendelian sampling and linkage. Genetics Research, 93(1):47–64.

Hoskins, R. A., Carlson, J. W., Wan, K. H., Park, S., Mendez, I., Galle, S. E., Booth, B. W., Pfeiffer, B. D., George, R. A., Svirskas, R., et al. (2015). The Release 6 reference sequence of the *Drosophila melanogaster* genome. Genome Research, 25(3):445–458.

Kardos, M., Luikart, G., and Allendorf, F. W. (2015). Measuring individual inbreeding in the age of genomics: marker-based measures are better than pedigrees. Heredity, 115(1):63–72.

Lenormand, T. and Dutheil, J. (2005). Recombination difference between sexes: a role for haploid selection. PLoS Biology, 3(3):e63.

Lian, J., Yin, Y., Oliver-Bonet, M., Liehr, T., Ko, E., Turek, P., Sun, F., and Martin, R. H. (2008). Variation in crossover interference levels on individual chromosomes from human males. Human Molecular Genetics, 17(17):2583–2594.

Mancera, E., Bourgon, R., Brozzi, A., Huber, W., and Steinmetz, L. M. (2008). High-resolution mapping of meiotic crossovers and non-crossovers in yeast. Nature, 454(7203):479–485.

Matute, D. R., Comeault, A. A., Earley, E., Serrato-Capuchina, A., Peede, D., Monroy-Eklund, A., Huang, W., Jones, C. D., Mackay, T. F., and Coyne, J. A. (2020). Rapid and predictable evolution of admixed populations between two *Drosophila* species pairs. Genetics, 214(1):211–230.

Sardell, J. M. and Kirkpatrick, M. (2020). Sex differences in the recombination landscape. American Naturalist, 195(2):361–379.

Sherman, P. W. (1979). Insect chromosome numbers and eusociality. The American Naturalist, 113(6):925–935.

Thompson, E. A. (2013). Identity by descent: variation in meiosis, across genomes, and in populations. Genetics, 194(2):301–326.

Turelli, M. and Orr, H. A. (2000). Dominance, epistasis and the genetics of postzygotic isolation. Genetics, 154(4):1663–1679.

Veller, C., Edelman, N. B., Muralidhar, P., and Nowak, M. A. (2019a). Recombination, variance in genetic relatedness, and selection against introgressed DNA. BioRxiv, page 846147. https://www.biorxiv.org/content/10.1101/846147v1.

Veller, C., Kleckner, N., and Nowak, M. A. (2019b). A rigorous measure of genome-wide genetic shuffling that takes into account crossover positions and Mendel’s second law. Proceedings of the National Academy of Sciences, 116(5):1659–1668.

Visscher, P. M., Macgregor, S., Benyamin, B., Zhu, G., Gordon, S., Medland, S., Hill, W. G., Hottenga, J.-J., Willemsen, G., Boomsma, D. I., et al. (2007). Genome partitioning of genetic variation for height from 11,214 sibling pairs. The American Journal of Human Genetics, 81(5):1104–1110.

Visscher, P. M., Medland, S. E., Ferreira, M. A. R., Morley, K. I., Zhu, G., Cornes, B. K., Montgomery, G. W., and Martin, N. G. (2006). Assumption-free estimation of heritability from genome-wide identity-by-descent sharing between full siblings. PLoS Genetics, 2(3):e41.

Wang, J. (2016). Pedigrees or markers: Which are better in estimating relatedness and inbreeding coefficient? Theoretical Population Biology, 107:4–13.

Wang, J., Fan, H. C., Behr, B., and Quake, S. R. (2012). Genome-wide single-cell analysis of recombination activity and de novo mutation rates in human sperm. Cell, 150(2):402–412.

Wang, S., Veller, C., Sun, F., Ruiz-Herrera, A., Shang, Y., Liu, H., Zickler, D., Chen, Z., Kleckner, N., and Zhang, L. (2019). Per-nucleus crossover covariation and implications for evolution. Cell, 177:326–338.

Weir, B. S., Avery, P. J., and Hill, W. G. (1980). Effect of mating structure on variation in inbreeding. Theoretical Population Biology, 18(3):396–429.

White, I. M. S. and Hill, W. G. (2020). Effect of heterogeneity in recombination rate on variation in realised relationship. Heredity, 124:28–36.

Wilfert, L., Gadau, J., and Schmid-Hempel, P. (2007). Variation in genomic recombination rates among animal taxa and the case of social insects. Heredity, 98(4):189–197.

Young, A. I., Frigge, M. L., Gudbjartsson, D. F., Thorleifsson, G., Bjornsdottir, G., Sulem, P., Masson, G., Thorsteinsdottir, U., Stefansson, K., and Kong, A. (2018). Relatedness disequilibrium regression estimates heritability without environmental bias. Nature Genetics, 50(9):1304–1310.

